# The Dynamics of Long Terminal Repeat Proliferation in the *Hesperis matronalis* Genome

**DOI:** 10.64898/2026.07.16.739001

**Authors:** Joanna L. Rifkin, Stephen E. Johnson, Arthur E. Weis, Stephen I. Wright, Regina S. Baucom

**Affiliations:** University of Toronto; University of Michigan

## Abstract

Genome sizes vary across four orders of magnitude in flowering plants, with consequences for evolution. Much of this variation is due to differences in transposable element content, particularly in long terminal repeat (LTR) retrotransposons. Despite their importance in plant genome evolution, LTRs have long been challenging to characterize because of their repetitive nature, but recent advances in sequencing allow more detailed explorations of their behavior. They are now known to occupy distinct genomic niches, and to evolve and proliferate over time as they escape host controls. In this study, we present a new genome sequence of the largest Brassicaceae genome, *Hesperis matronalis*, and describe its transposable element complement and how LTRs contributed to its genome expansion. We find evidence for both early proliferation of Ty3 elements and rapid recent expansion of Ty1-copia elements. In addition, we place the *H. matronalis* LTRs in a broader context of LTR evolution in the Brassicaceae, showing that the dominant copia families are part of an evolutionary radiation endemic to *Hesperis*. Finally, we describe differences in LTR age, proximity to genes, and apparent removal rate which suggest consistent genomic niches over the lifespan of a TE family. These results shed light on how LTRs evolve dynamically with host genomes, and have contributed to the expansion of the largest genome in the Brassicaceae.

**Significance Statement:** Organisms differ in many ways, including the size of their genomes. Genome size, in turn, can affect many aspects of evolution, such as what kinds of mutations are available to natural selection and how effectively natural selection can act. The “complexity” of an organism does not predict its genome size; rather, much of this variation is explained by the amount of transposable elements, or “jumping genes,” that inhabit a genome. This study describes the evolutionary history of the transposable elements in dame’s rocket (*Hesperis matronalis*), which has the largest genome of any plant in the mustard family (Brassicaceae). We find that the genome inflation in *Hesperis* is due to transposable elements that are not found elsewhere in the Brassicaceae, and likely diversified within *Hesperis*. Characterizing the evolution and behavior of transposable elements in this genome offers a better understanding of the forces that determine genome size and ultimately affect the evolution of life on earth.

## Introduction

Generations of biologists have dedicated themselves to understanding the diversity of life on earth - Darwin’s oft-quoted “endless forms most beautiful” (Darwin 1859) - and improvements in both theoretical frameworks and measurement technologies reveal ever more axes of variability. After DNA was identified as the heritable material, estimates of DNA content revealed that genome size varied widely but did not scale with organismal “complexity” (Thomas Jr 1971), an observation dubbed the “C-value paradox.” Angiosperms in particular have hugely variable genome sizes, ranging from under 100 megabases (Mb) to approximately 150,000 Mb (Elliott and Gregory 2015; Pellicer et al. 2018). As complete sequences have become available for more genomes, it has become clear that gene number alone does not explain this variation; in angiosperms, for example, the smallest genomes have between 10,000 and 20,000 genes (Vu et al. 2015), while the largest have somewhat over 100,000 (Jiao et al. 2025). While gene number does increase with genome size, it is inadequate to explain the variation. Instead, angiosperm genome sizes appear to expand through whole genome duplication (Van de Peer et al. 2017; Clark and Donoghue 2018) and, particularly, the proliferation of repetitive sequence (Kubis et al. 1998; Hawkins et al. 2009; Ågren and Wright 2011; Macas et al. 2015; Gaiero et al. 2019; Wells and Feschotte 2020); in a large-scale comparative analysis, genome size variation is best explained by the relative proportions of genic and repetitive sequence (Elliott and Gregory 2015).

Genome size evolution is dynamic, and genome size itself, in turn, shapes other aspects of biology and evolution. On an organismal level, genome size can affect plant size, local abundance, and competitive ability (Herben et al. 2012). Across evolutionary time, it has been suggested that larger, more repetitive genomes experience more structural rearrangements (Yu et al. 2011), a higher proportion of regulatory relative to coding mutations, and a higher prevalence of soft relative to hard selective sweeps (Mei et al. 2018). Once expanded, large genomes can also contract, chiefly through ectopic recombination eliminating blocks of repetitive sequence (Hawkins et al. 2009). Genome size and content are thus consequential forces essential to understanding plant evolution (Lisch 2013; Mei et al. 2018). However, our knowledge of the relative importance of these processes and how they interact remains limited by the difficulty of resolving repetitive sequence, particularly in larger genomes (Warburton and Sebra 2023). Recent advances in sequencing technology and the increasing availability of the sequences of more diverse plant genomes (Sun et al. 2022) offer the promise of greater insight into the role of repetitive sequence in genome and genome size evolution.

The dominant type of repetitive sequence in angiosperm genomes is transposable elements (TEs), particularly retrovirus-related long terminal repeat retrotransposons (LTRs) (Kubis et al. 1998; Hawkins et al. 2009; Ågren and Wright 2011; Macas et al. 2015; Gaiero et al. 2019; Wells and Feschotte 2020). Transposable elements can be understood to inhabit the genome as an “ecosystem,” occupying distinct “niches” (Brookfield 2005; Venner et al. 2009; Sultana et al. 2017; Stitzer et al. 2021) and exhibiting evolutionary dynamics, which are particularly well characterized for LTRs. TEs interact in various ways with the host genome and its evolution (reviewed in (Lisch 2013; Schrader and Schmitz 2019; Catlin and Josephs 2022)): transposable elements in and near genes can have direct effects on phenotype through disrupting coding sequence and through spillover epigenetic silencing (Lee and Karpen 2017), and by increasing the likelihood of ectopic recombination that contributes to structural rearrangements (Berdan et al. 2023). LTRs are generally more abundant in gene-poor regions (Baucom et al. 2009a; Stitzer et al. 2021) and exhibit diverse proliferation patterns, with some evidence for both stochastic bursts and slower, steadier accumulation often occurring simultaneously in different TE lineages within the same host genome (Dai et al. 2018). TE bursts have been suggested to be associated with changes to the genomic environment, including stress (Belyayev 2014; Potapenko et al. 2026), while patterns of LTR abundance and distribution may also reflect selection acting on LTR families (Baucom et al. 2009b). Because LTRs can be both classified based on coding sequence and approximately dated using a mutational clock, both current and historical variation in abundance, proliferation rates, and removal can be observed at multiple levels: for example, different LTR families proliferated at different times between the subgenomes in polyploid cotton, teff, and strawberries (Lyu et al. 2025), between tropical and temperate maize accessions (Ou et al. 2024), between cotton species (Hawkins et al. 2009), across the genus *Solanum* (Gaiero et al. 2019), and throughout the *Fabeae* tribe (Macas et al. 2015).

Despite the well-documented variation and evolutionary dynamics of LTRs, however, their evolutionary trajectories and relationship to the genomic environment remain unpredictable. Much of the variation in age and abundance of LTRs appears to be family-specific and not explained by features of either genomic region or host identity (Stitzer et al. 2021; Ou et al. 2024). In addition, the most comprehensive descriptions of LTR ecology so far have been largely focused within maize (Baucom et al. 2009a; Stitzer et al. 2021; Ou et al. 2024), and while maize hosts an astonishing diversity of TEs and impressive genome size variation for one species (Bilinski et al. 2018), its TE dynamics may not be universally representative.

Therefore, reaching a more complete understanding of the community ecology of transposable elements requires in-depth characterizations of TEs across a range of genome sizes. This variation is available within the Brassicaceae (Mandáková et al. 2017). Most Brassicaceae have small genomes, which comparative approaches suggest have been stable across a long span of evolutionary time (Lysak et al. 2009). In contrast, species in the genus *Hesperis* have experienced extensive genome inflation. *Hesperis matronalis*, a weedy species with both diploid and polyploid populations widespread in Eurasia and invasive in North America, boasts the largest single-copy genome within the family (Francis et al. 2009; Hloušková et al. 2019; Eslami-Farouji et al. 2021). Low-pass sequence data and modeling indicate a recent burst of Ty1-Copia elements likely expanded *Hesperis* genomes, with earlier Ty3 (formerly Gypsy) expansion also contributing substantially to inflated genomes within the clade (Hloušková et al. 2019; Beric et al. 2021).

However, short read sequencing offers limited ability to resolve repetitive sequences, particularly at a large scale (Warburton and Sebra 2023), and fluorescent cytology is less tractable in extremely large genomes (Hloušková et al. 2019). Therefore details of the timing and extent of LTR proliferation within *Hesperis* remain unknown, particularly the relative timing of Copia and Ty3 expansion and their positioning within the genome, which lineages within these superfamilies are most responsible for the genome expansion, and the extent of within-*Hesperis* diversification compared to the diversification of these elements across related species. TEs can proliferate both gradually and in transient, stochastic bursts dominated by only a small subset of TE lineages (El Baidouri and Panaud 2013; Ou et al. 2024; Liu et al. 2025). Rapid TE expansions may co-occur with environmental stresses and speciation events (Belyayev 2014). However, expansion dynamics vary widely across both TE and host lineages and across evolutionary time (Hawkins et al. 2009; Dai et al. 2018; Ou et al. 2024), and while transposable elements appear to be responsible for genome size expansion in *H. matronalis* (Lysak et al. 2009; Mandáková et al. 2017; Hloušková et al. 2019), it remains unclear how this expansion proceeded. Characterizing the TE landscape and dynamics of an unusually inflated genome will shed light on these dynamics in an extreme case.

To this end, we present a new *de novo* assembly of a diploid accession of *Hesperis matronalis* in combination with a gene annotation and a pan-TE annotation across major clades in the Brassicaceae to investigate the dynamics of a major, sustained LTR burst in its broader evolutionary context. With these resources, we explore the following questions: (1) what were the relative timings of Ty3 and copia expansion in the *H. matronalis* genome? (2) To what extent is LTR burden conserved across populations within *H. matronalis*, and does a diploid reference genome capture species-wide patterns of LTR abundance? (3) Did LTR expansion occur broadly across Ty3 and copia clades, suggesting a general change to the genomic environment, or concentrate instead within a few successful lineages, consistent with family-specific escape from host repression? (4) Have genomic niches remained stable across TE expansion events, or do LTR proliferation events coincide with the invasion of new niches?

## Results and Discussion

### Genome size estimates, assembly size and statistics

Existing karyotypes and flow cytometry estimates of the *H. matronalis* genome size indicate that both diploid and tetraploid populations occur, with 2N=14, 16, 24, 26, 28 all reported (Francis et al. 2009), and a 1C genome size of 3.72 Gb (Kubešová et al. 2010). We collected flow cytometry genome size estimates from 33 populations from the midwestern United States and southern Canada (Table S1). Our flow cytometry estimates, standardized relative to *Monstera deliciosa* and *Clivia miniata*, were mostly consistent with tetraploid genomes with a 3.6-4.3 Gb single-copy genome size (14.8-17.4 pg; Table S2). However one site (RIGA, lat. 41.816, long. -83.824) contained eight apparently diploid samples (7.3-7.8 pg) and two samples with peaks at both diploid and tetraploid sizes (7.36-8.04 and 14.5-15.8), suggesting either mixoploidy or endopolyploidy. We chose a single apparently diploid plant from this site to assemble a reference genome. The final filtered assembly totaled 3.87 Gb, consistent with flow cytometry estimates of haploid genome size for individuals from the source population, which ranged from 3.62 Gb to 4.02 Gb depending on flow cytometry standard (Table S2, Table S3). Assembly contiguity was high (N50 = 425.8 Mb), and completeness was estimated at 99.61% based on recovery of embryophyte BUSCO (Seppey et al. 2019) genes. The GC content of the genome was 41.55%. We recovered ten major scaffolds, with an expected N of six or seven (Francis et al. 2009), and performed most analyses on either all scaffolds over 100Mb (N=10, 3.35 Gb) or all scaffolds over 10Mb (N=14, 3.48 Gb).

Our BRAKER3 (Gabriel et al. 2024) annotation identified 43,313 gene models with a mean length of 357.4 amino acids (3222.55 base pairs) and a mean exon number of 4.1 (range 1-79). When we removed genes with any exons that overlapped an annotated transposable element (see below), 30,920 remained (Table S4, Table S5). Although we did not conclusively annotate centromeres, we used TRASH (Wlodzimierz et al. 2023) to identify satellite sequences that ranged in length from 7bp (consistent with the canonical plant telomere TTTAGGG; (Peska and Garcia 2020)) to 318bp and in frequency from 4285-104954 occurrences genomewide. Five satellites exhibited strongly localized peaks on one or more scaffolds (Fig. S1), and two were highly similar to satellites of identical lengths found using low-pass sequencing and validated with fluorescence in-situ hybridization in *H. sylvestris* (satellite 161_4, 93% identity with HeSy2; 174_1, 94% identity with HeSy5 (Hloušková et al. 2019); Table S6). None of the satellite sequences identified shared significant BLAST identity with any sequences in the NCBI database.

### Ty3 and copia elements sequentially expanded the *H. matronalis* genome

We annotated transposable elements in *Hesperis matronalis* by generating a panEDTA library including eleven species with long-read genome assemblies spread across the Brassicaceae (Table S7). The *H. matronalis* repeatome included helitrons, TIRs, SINEs, LINES, but as in many angiosperms, long terminal repeats (LTR) retrotransposons dominates the repeat landscape, comprising approximately 67% of sequence content on scaffolds over 50Mb, while DNA transposons and non-LTR retrotransposons combined contributed approximately 5% of sequence content (Table 1; Table S8). The majority of identifiable LTRs were Ty3 (20% of total sequence; 912958 total elements with 45827 intact) followed by copia (9% of total sequence; total elements 588551 with 47887 intact). The two transposable element superfamilies differed in their distribution across the genome: copia elements were more evenly distributed across chromosomes, while Ty3s were concentrated in gene-poor regions possibly near centromeres (Fig. 1). Consistent with this, copia elements were significantly closer to genes than Ty3s (mean 137155.1bp vs. 151389.1bp, median 82284.5bp vs. 97392.5bp, t statistic -40.261 with 729575 df, p < 2.2e-16; Test 1 in analysis script).

**Fig. 1.**
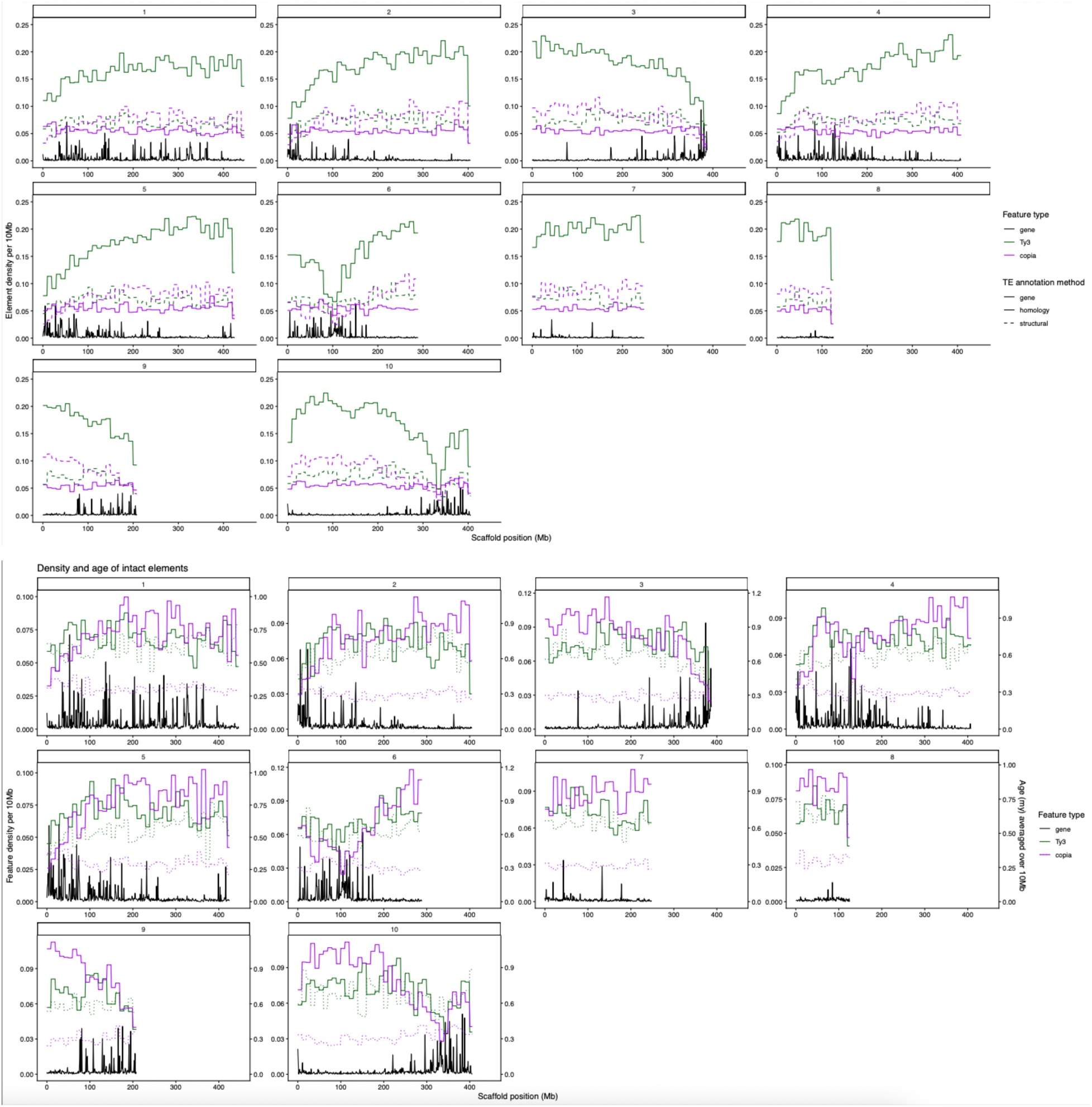
Landscape of genes and transposable elements in the *H. matronalis* genome in ten-megabase bins. Top: Panels each represent one major scaffold, color indicates feature type (black: gene; green: Ty3 element; purple: copia element) and line type distinguishes intact structurally annotated (dashed) from fragmented homology-annotated (solid) transposable elements. Bottom: Panels each represent one major scaffold, color indicates feature type of intact elements only (black: gene; green: Ty3 element; purple: copia element), dotted line indicates age of intact LTRs (right axis). Alt text: This figure shows two ten-panel displays with graphs of the distributions of genes and two types of long terminal repeat retrotransposons along the scaffolds of the draft genome assembly.

**Table 1:** Repetitive elements in the *H. matronalis* genome.

|  | Number of elements* | Length occupied | Percent of sequence |
| --- | --- | --- | --- |
| Retroelements | 3323954 | 2215836612 | 66.567 |
| SINES | 142411 | 13833819 | 0.396 |
| Penelope | 0 | 0 | 0 |
| LINEs | 98038 | 39076760 | 1.149 |
| CRE/SLACS | 0 | 0 | 0 |
| L2/CR1/Rex | 6181 | 1358217 | 0.039 |
| R1/LOA/Jockey | 0 | 0 | 0 |
| R2/R4/NeSL | 0 | 0 | 0 |
| RTE/Bov-B | 0 | 0 | 0 |
| L1/CIN4 | 91857 | 37718543 | 1.109 |
| LTR elements | 3083505 | 2162926033 | 65.022 |
| BEL/Pao | 14 | 7756 | <0.001 |
| Ty1/Copia | 588551 | 314694276 | 9.414 |
| Ty3/DIRS1** | 912958 | 677748459 | 20.42 |
| Retroviral | 0 | 0 | 0 |
| DNA transposons | 311408 | 122547055 | 3.614 |
| hobo-Activator | 0 | 0 | 0 |
| Tc1-IS630-Pogo | 0 | 0 | 0 |
| En-Spm | 0 | 0 | 0 |
| MuDR-IS905 | 0 | 0 | 0 |
| PiggyBac | 0 | 0 | 0 |
| Tourist/Harbinger | 0 | 0 | 0 |
| Other | 0 | 0 | 0 |
| Rolling-circles | 0 | 0 | 0 |
| Unclassified | 790849 | 463914586 | 13.716 |
| Simple repeats | 224381 | 11477762 | 0.338 |
| Low complexity | 48416 | 2640420 | 0.078 |
| Small RNA | 0 | 0 | 0 |
| Satellites | 0 | 0 | 0 |
| Total interspersed repeats |  | 2802298253 | 83.896 |

\* Nested elements may be counted as one by EDTA.
\*\* Formerly "Gypsy."

Previous investigations in the Brassicaceae have implicated both Ty3 and copia LTRs in the large size of the *Hesperis matronalis* genome, with a particular role for a recent copia expansion (Beric et al. 2021). To explore this, we investigated the age and proliferation rates of the two major superfamilies of LTRs. Because the long terminal repeats that define these transposable elements lose identity when inactive, structurally annotated intact LTRs can be assumed to have inserted more recently than incomplete LTRs identified by homology. As would be expected if copia LTRs are younger, 34.9% of copia elements in the *H. matronalis* genome are intact, compared with 12.6% of Ty3 (chi sq with one df, n = 19805, p < 2.2 e-16; Fig. 1; Test 2 in analysis script). LTR age can also be estimated based on divergence between the terminal repeats using a mutational clock; this approach also supported a younger average for copia elements (296422.6 years, compared to 620040.8 years for Ty3 elements; t statistic - 92.875 with 37344 df, p < 2.2e-16; Test 3 in analysis script). The distribution of insertion ages of intact copia elements further supports more recent proliferation than Ty3 elements, both across all (Fig. 2a) and the most numerous (Fig. 2b) elements (Table S9). Note that these distributions likely reflect a combination of not only differences in the timing of transposition but also differences in the strength of natural selection; since copia elements tend to be closer to genes, they will be more likely to be subject to negative selection against insertions, driving their age distribution towards the younger end.

**Fig. 2.**
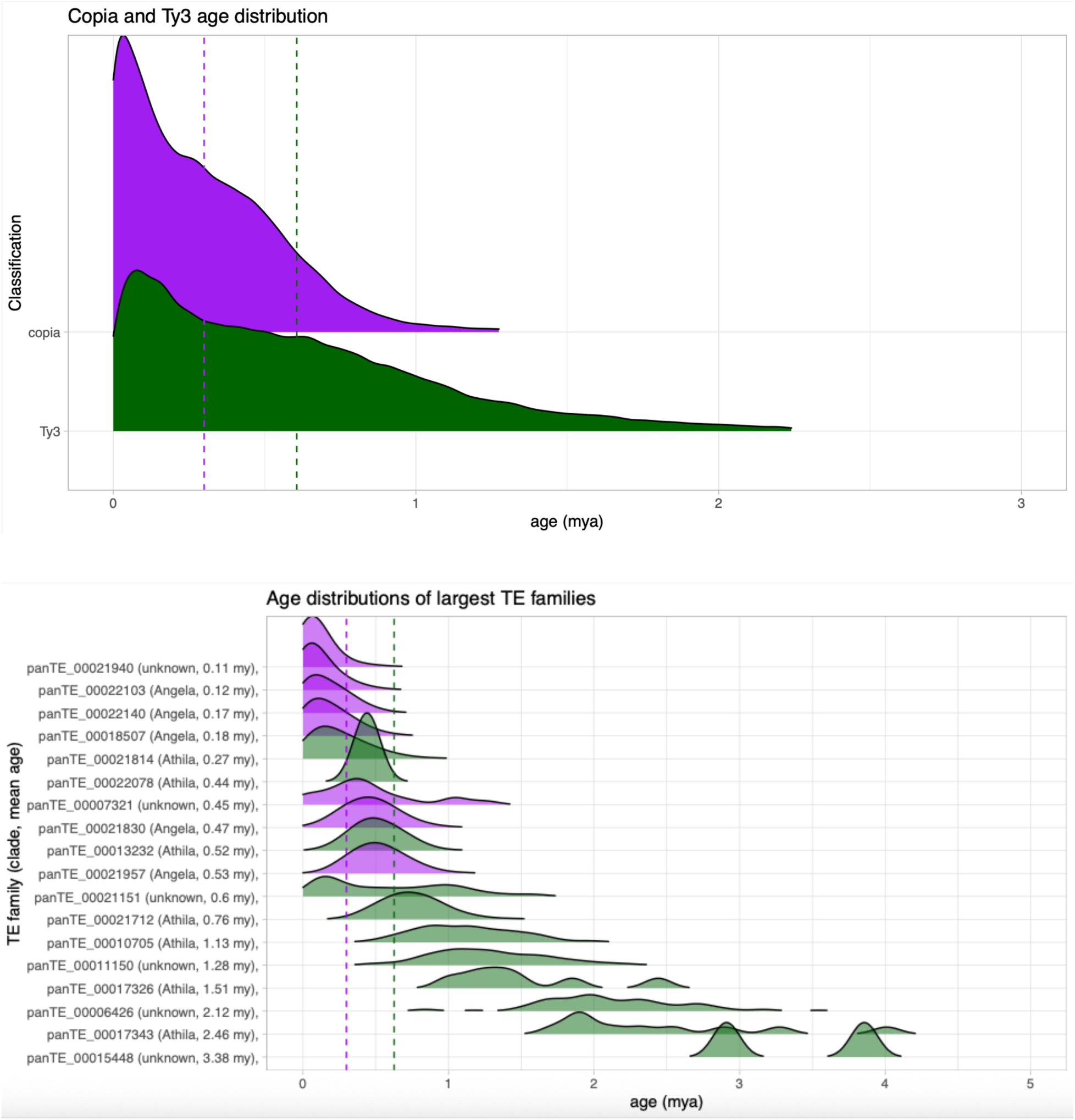
Age distributions of all (top) and most populous (bottom) intact copia (purple) and Ty3 (green) elements in the *Hesperis matronalis* genome. Dashed lines indicate mean ages of all intact copia (purple) and Ty3 (green) elements. Clade and mean family age in parentheses. Alt text: This figure shows two one-panel displays with ridge plots of age distributions of transposable element families.

Finally, we roughly estimated the relative proportions of copia and Ty3 elements from restriction site-associated DNA sequencing (RADseq; (Peterson et al. 2012)) samples across 33 North American populations using the TERad pipeline (Chak and Rubenstein 2019). Even with the reduction in genome coverage, these estimates were strikingly consistent both across populations and with the proportions observed in whole-genome analysis. In the sequenced *H. matronalis* genome, LTRs accounted for a total of 64% of total genomic content, with Ty3 accounting for 20% and copia for 9%. An estimate from RADseq from the population where the sequenced individual was collected estimated total LTRs as 54.7% of genome content, copia elements as 9.4%, and Ty3 as 34.2%. The higher estimate for Ty3 may reflect the age and diversity of elements that can be more accurately classified with complete sequence across repetitive regions, along with possible sequencing biases from RADseq. Across the 47 populations surveyed, these proportions ranged from 48%-61.6% total, 6.3%-10.8% copia, and 31.4%-38.8% Ty3, all of which are broadly consistent with the estimate from the genome annotation. Further, RIGA, the source population of the sequenced individual, was not an obvious outlier in the distribution (Fig. 3; Table S10). We can thus be confident that although our sequenced accession is diploid, it broadly represents TE burden of the species, and, consistent with previous research (Hloušková et al. 2019), supports the idea that the expansion of the *H. matronalis* genome via LTR proliferation likely predated genome duplication.

**Fig. 3.**
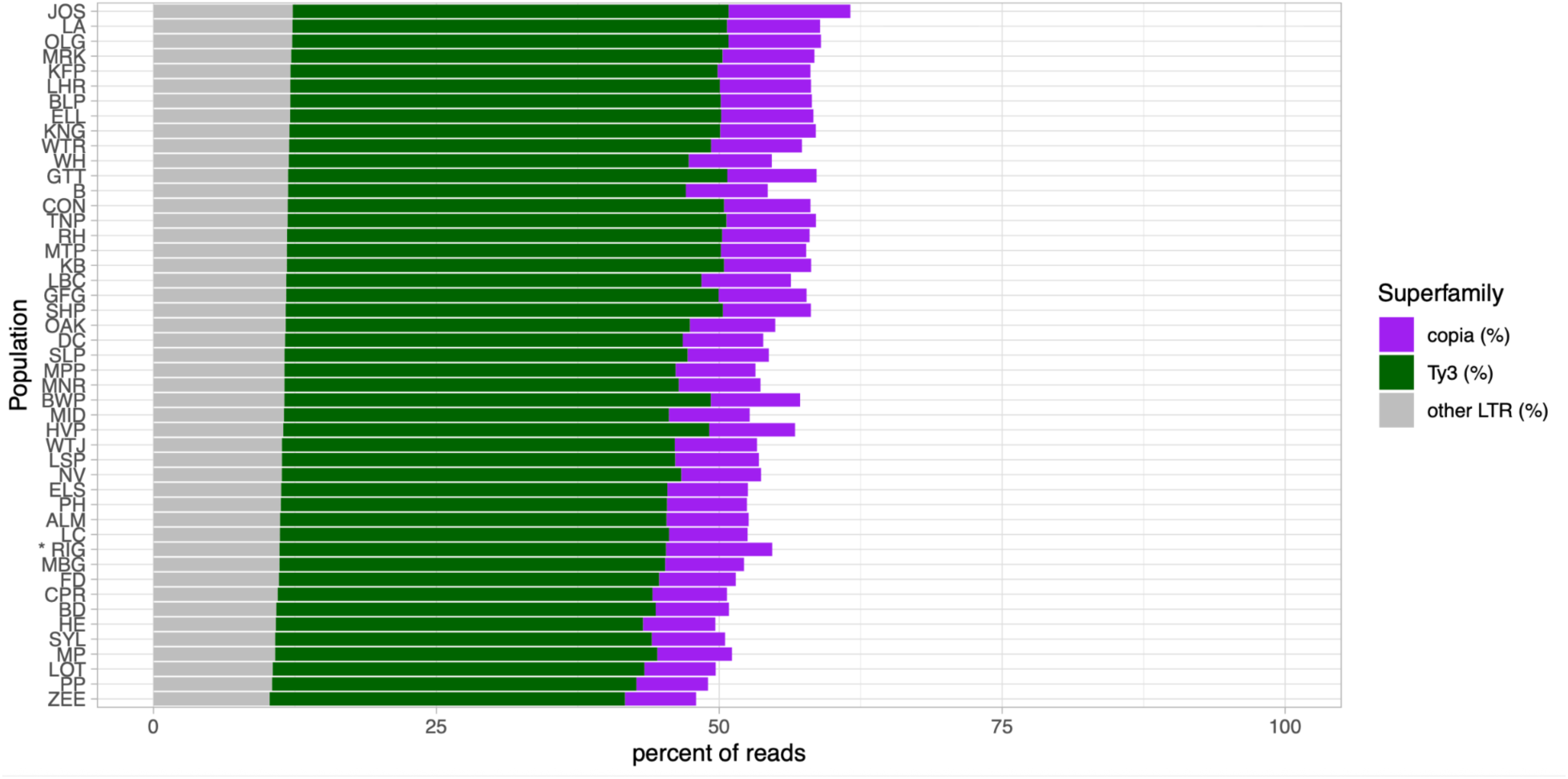
Percentage of LTRs in North American genomes, estimated from ddRAD data. Purple indicates copia, green indicates Ty3, gray indicates other LTRs. The population from which the sequenced genome was sampled (RIG) is indicated with an asterisk. Alt text: This figure shows a barplot of the amount of long terminal repeat retrotransposons across multiple populations.

### Comparative genomic analysis reveal extensive copia diversification and turnover

Our superfamily-wide results are consistent with multiple waves of genome expansion in the *H. matronalis* lineage. Recent copia expansion appears to have contributed to recent genome size increases within *Hesperis matronalis*, while earlier and ongoing Ty3 proliferation likely increased genome sizes across a longer evolutionary timescale. To better understand this timescale, we next placed this expansion into a family-wide context across the Brassicaceae. *H. matronalis* has more than twice as many TEs in relative terms than any other species examined, and over an order of magnitude more in absolute terms (Table 2; as in *H. matronalis*, non-LTR retrotransposons and DNA transposons did not constitute more than 5% of any genome). Consistent with a recent copia expansion, *H. matronalis* also had a higher genomic proportion of copia elements than any other genome examined, and they were on average younger than other species in the Hesperideae and among the youngest in the Brassicaceae (Fig. 5, Table S11). Ty3 abundance was also considerably increased compared to other species, but with less evidence for very recent expansion (Fig. 5).

**Fig. 4.**
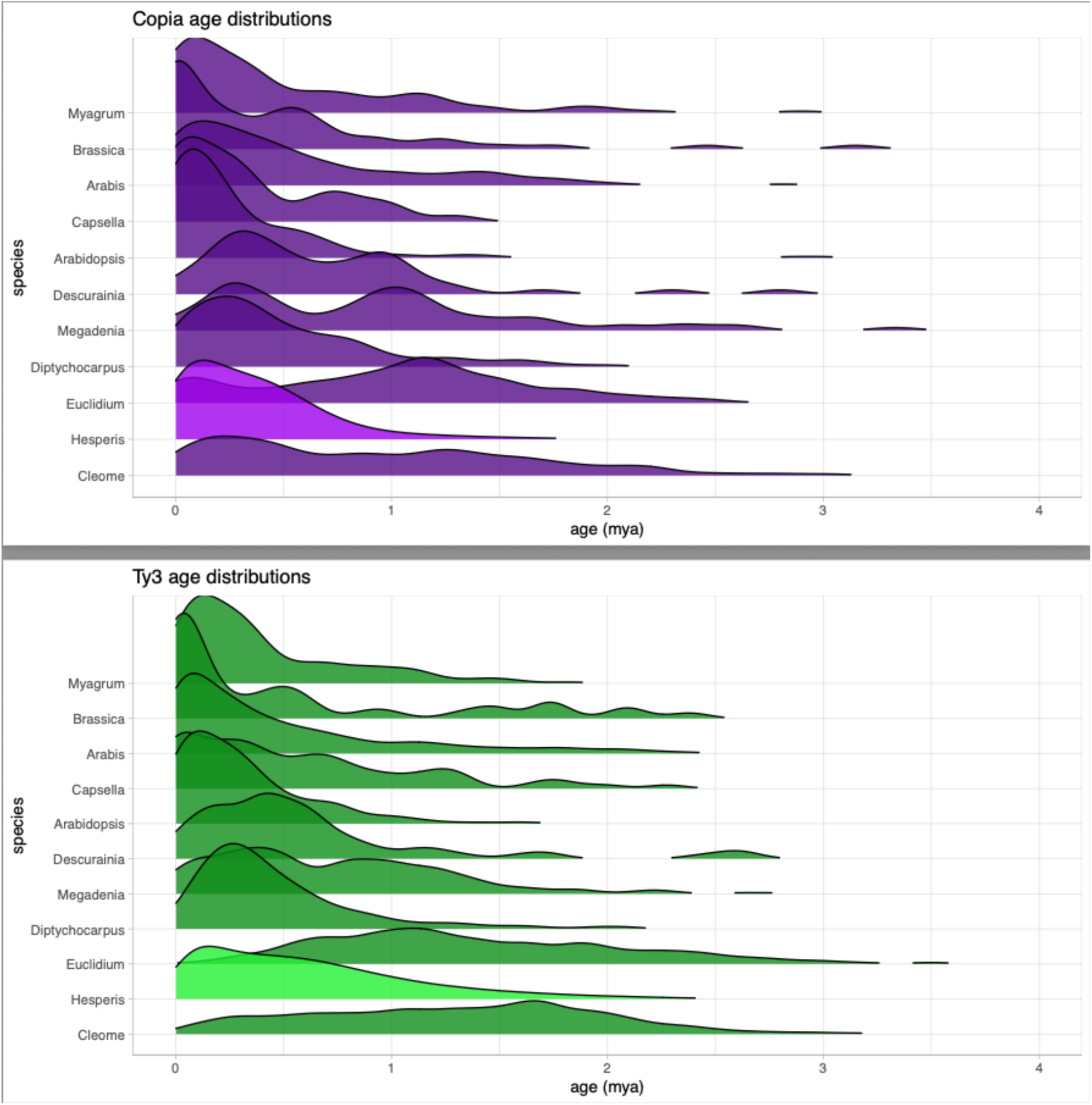
Age distributions of intact transposable elements across species. Alt text: This figure shows two one-panel displays with ridge plots of age distributions of transposable elements in different species in the Brassicaceae.

**Fig. 5.**
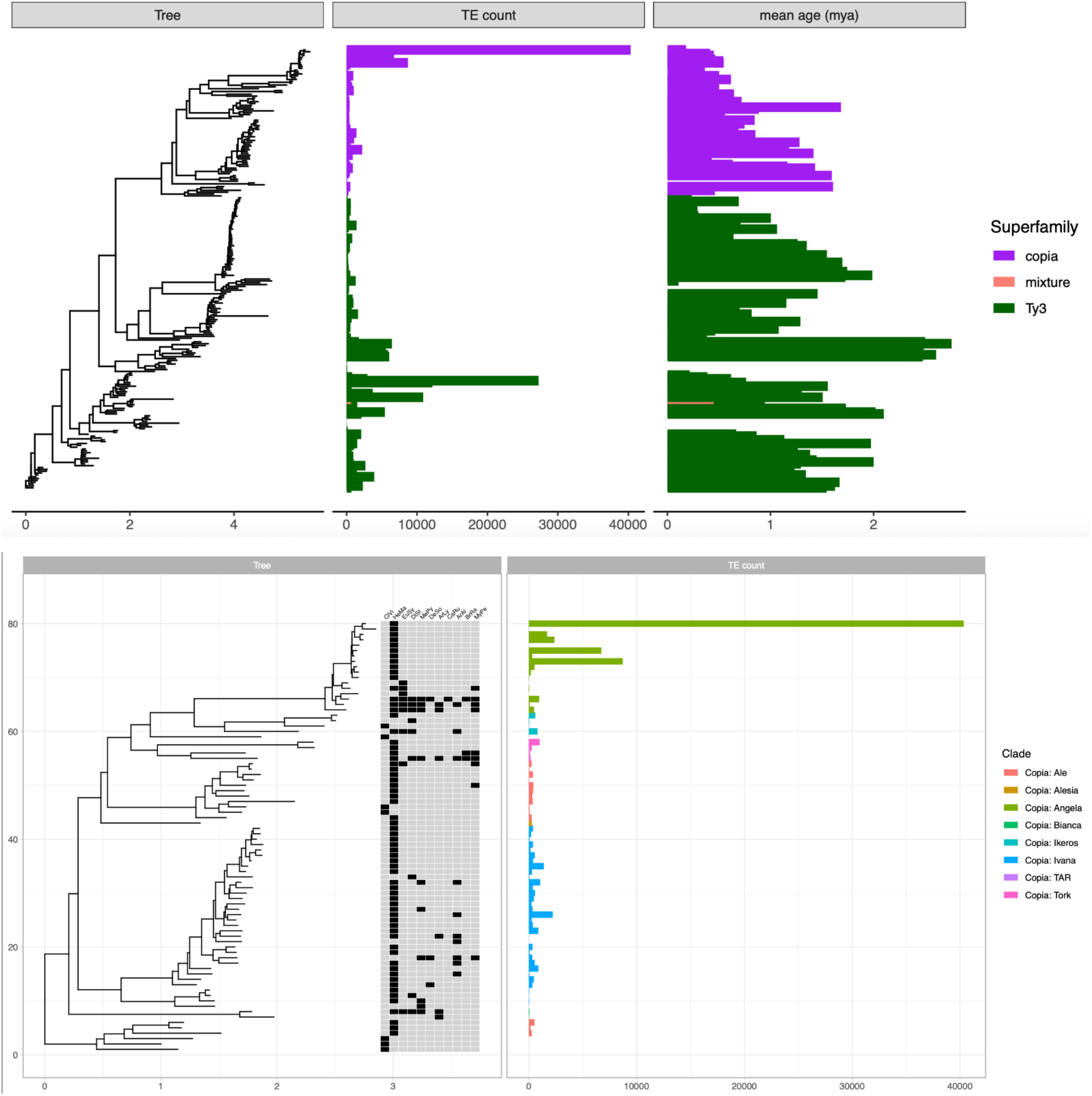
Top: Phylogeny of all identified Brassicaceae copia and Ty3 LTRs with TE counts and ages. Bottom: phylogeny of copia elements with clades and matrix of presence/absence across Brassicaceae. Alt text: This figure shows two one-panel displays with phylogenetic trees of transposable elements, their ages, their frequency, and their taxonomic distribution across the Brassicaceae.

**Table 2:**
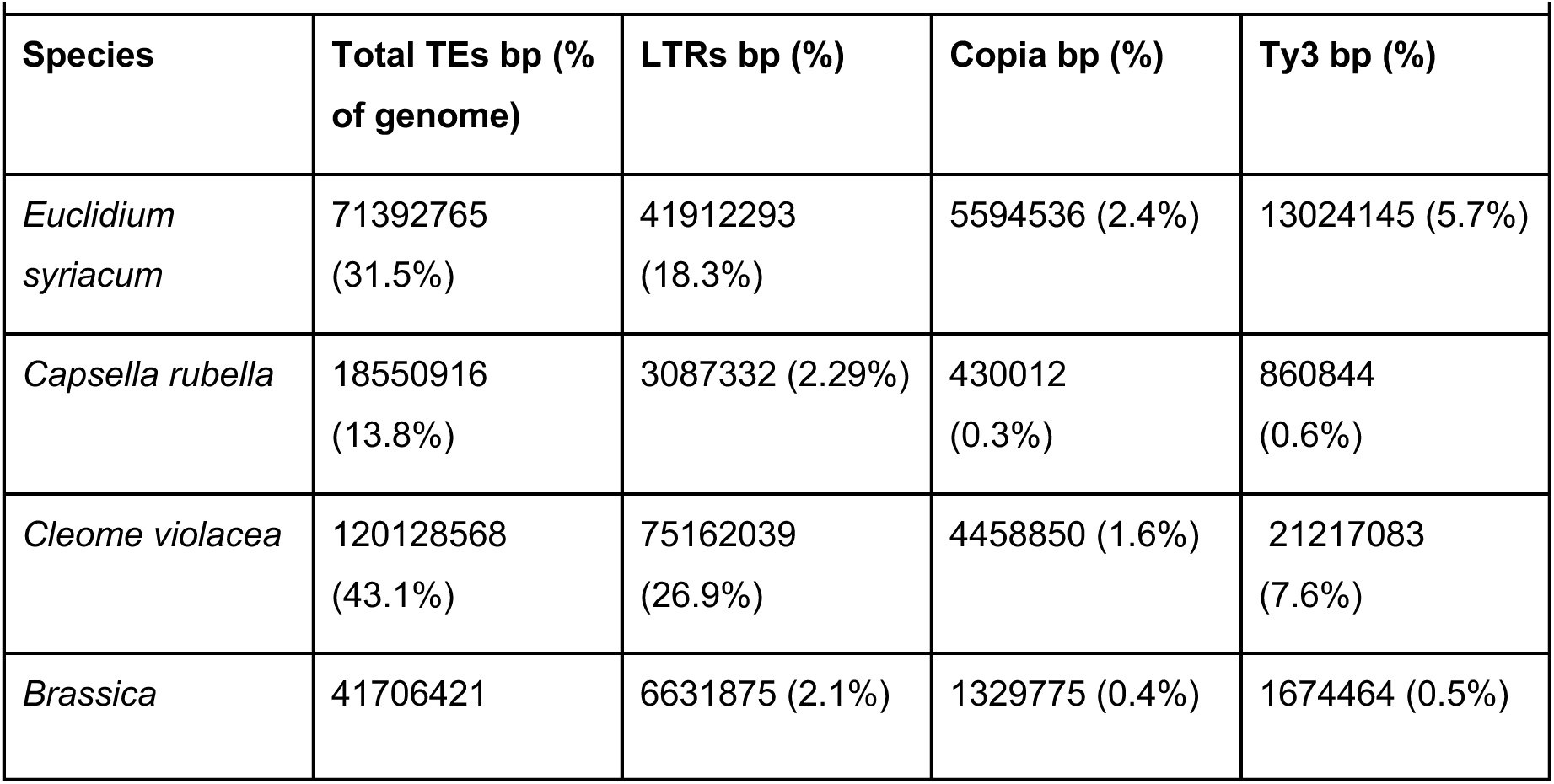
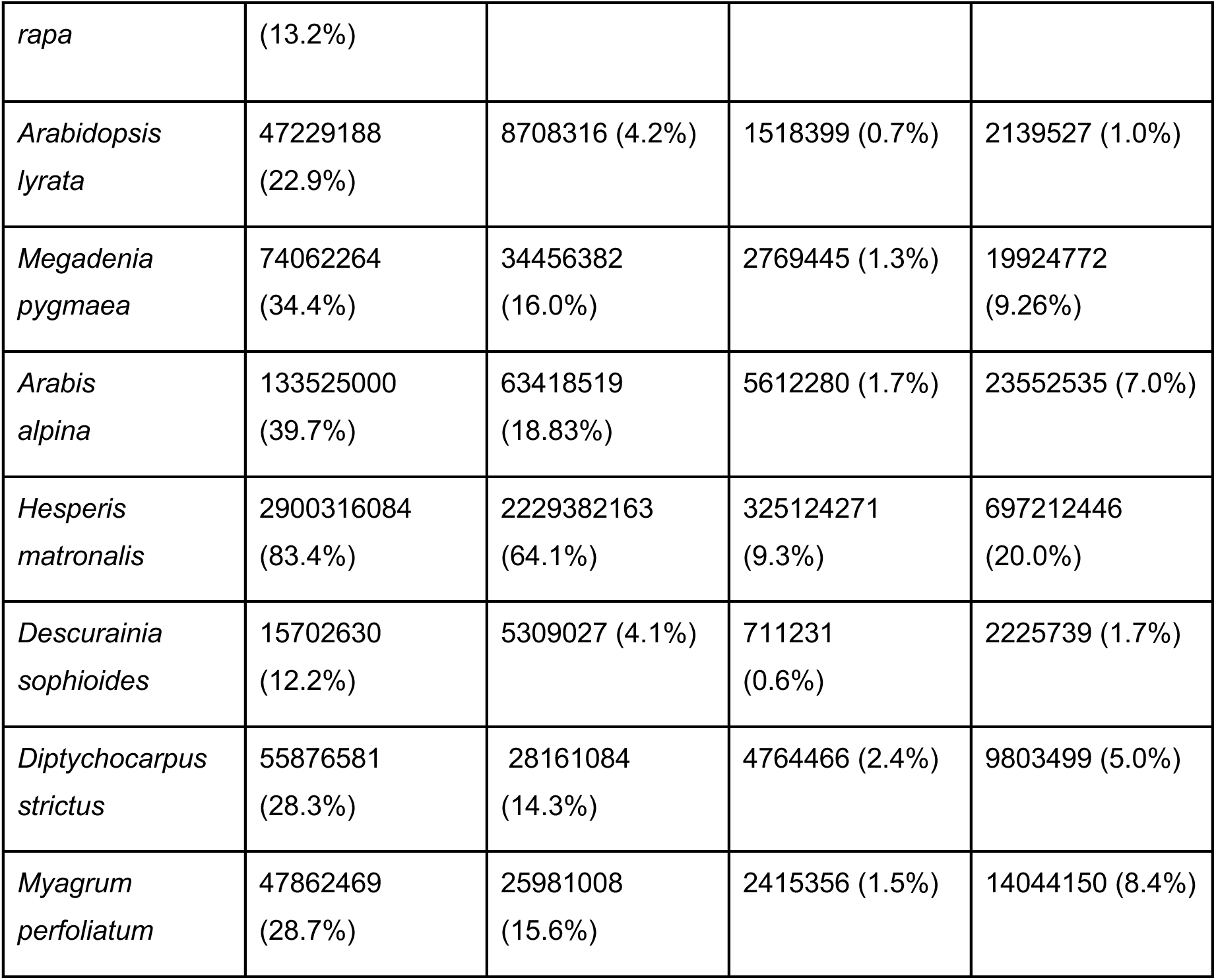
Comparative transposable element burden in Brassicaceae bp (%)

We then explored the roles of different lineages within these superfamilies by using TESorter (Zhang et al. 2022) to classify all LTRs that were full length in at least one genome in our family-wide panEDTA library. 2554 sequences from our library could be categorized, of which 2443 occurred in *Hesperis matronalis*; the most diverse clades were the copia taxa Angela and Ale, and the Ty3 taxa Athila, CRM, and Retand (Table S12). We constructed a phylogeny LTRs using the GAG/PROT genes sequences extracted by TESorter. Within *H. matronalis*, the largest and youngest families were concentrated within the Angela lineage in a group of 11 LTR families unique to *H. matronalis*, sister to a branch containing a family unique to *Euclidium syriacum* and a family shared between *E. syriacum*, *H. matronalis*, and *Myagrum perfoliatum*. This is consistent with the copia expansion in *H. matronalis* being largely driven by local diversification of a *Hesperis*-specific Angela lineage. In LTR expansions in other species, the pattern of dominance by one or a few families is widespread (El Baidouri and Panaud 2013).

In a comparative context, transposable elements may also facilitate ectopic recombination and contribute to genomic rearrangements and TE removal. Because of its heavy TE burden, *H. matronalis* might be expected to exhibit dramatic rearrangements relative to related species. To describe the extent of rearrangements in *H. matronalis*, we performed a synteny analysis using GENESPACE (Lovell et al. 2022). Despite the massive genome expansion of *H. matronalis*, it appears to retain considerable synteny to other species in the Brassicaceae, particularly compared to a highly rearranged genome like *B. rapa* (Fig. 6). Although our assembly likely does not fully recapitulate complete chromosomes, large-scale syntenic features are consistent with BAC-based reconstructions of ancestral Brassicaceae karyotypes (Mandáková et al. 2017). For example, *H. matronalis* chromosome 6 comprises homologs of *A. lyrata* chromosomes 4 and 5, whereas chromosome 3 is composed primarily of sequence homologous to chromosome 8 with an inserted region derived from chromosome 6. These concordant patterns suggest that, despite incomplete scaffold joining, the assembly is broadly reliable.

**Fig. 6.**
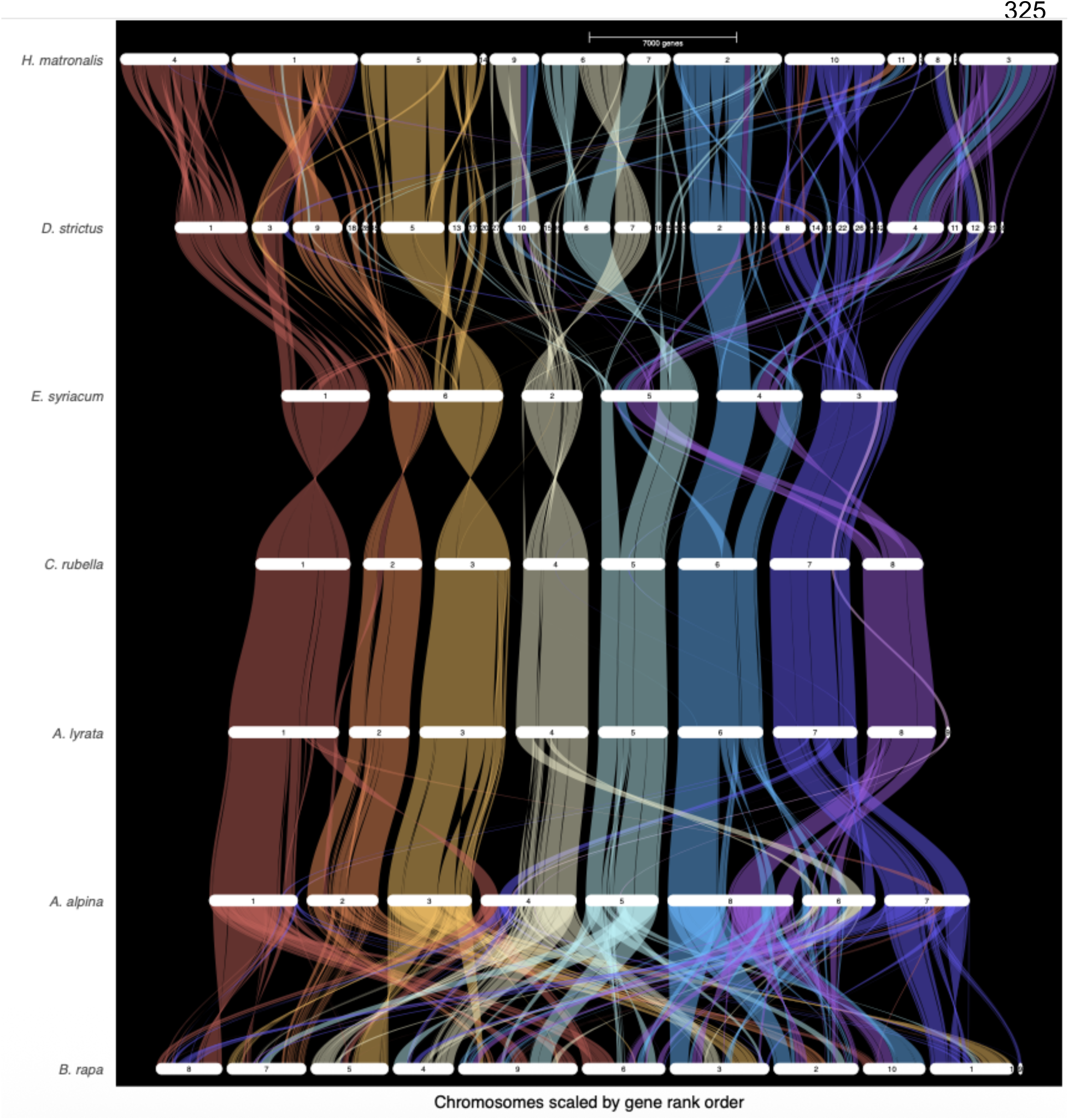
Gene-level synteny between *H. matronalis* and other Brassicaceae. Colored ribbons indicate syntenic blocks of genes relative to *A. lyrata* scaffolds. Species: *Hesperis matronalis*, *Diptychocarpus strictus*, *Euclidium syriacum*, *Capsella rubella*, *Arabidopsis lyrata*, *Arabis alpina*, *Brassica rapa*. Alt text: This figure shows a riparian plot of gene-level synteny across Brassicaceae species.

### No evidence of expanded niche for expanded LTRs

Typically, and across our dataset, Ty3 LTRs are strongly associated with pericentromeric regions, while copia LTRs are found both in genic and pericentromeric regions. These genomic “niches” have been observed across multiple taxa (Brookfield 2005; Venner et al. 2009; Stitzer et al. 2021), but it is unclear how stable they are across evolutionary time, and whether these dynamics shift during a TE expansion. To determine how niches shift across expansions and suppression, we investigated whether proliferating LTR families inhabit different genomic niches than families that proliferated less recently.

First, to identify contributors to the ongoing expansion, we identified the peaks of age distributions across copia families and, following (Ou et al. 2024), classified them into “young,” “moderate,” and “old” families. “Young” families that are in an ongoing expansion were defined as families in which the peak of the family’s age distribution overlaps zero (terminal repeats are identical), while “moderate” families had a peak that did not overlap zero but included at least 5% intact copies with identical terminal repeats, and “old” families were those in which less than 5% of LTRs had identical terminal repeats (Table 3). By these criteria, the distributions of family ages in *H. matronalis* Ty3 and copia do not differ (table; G statistic 4.3522 with 2 df, p-value = 0.1135; Test 4 in analysis script). However, young copia families were significantly larger than young Ty3 families (copia: mean 2649.024 copies, n=41 families; Ty3, mean 66.1 copia n=225 families; t statistic 2.0481 with 40.004 df, p =0.04715; Test 5 in analysis script); we also identified the largest families overall (those with over 10000 representatives), and found that while 4 of the 7 copia families in this group were young (moderate: 1; old: 2), all 10 of the largest Ty3 families were either moderate (2) or old (8).

**Table 3:** **Distribution of family ages in Ty3 and copia within *H. matronalis***

|  | Young | Moderate | Old |
| --- | --- | --- | --- |
| Ty3 | 225 | 117 | 1159 |
| Copia | 41 | 26 | 166 |

We next explored the interplay between proliferation and removal in LTR age distributions. If both superfamilies are proliferating rapidly but copia elements are removed more frequently, we predict that fewer older copia elements will remain, i.e., that copia elements will on average be younger. Alternatively, if copia elements are both proliferating rapidly and removed rapidly, the current landscape may reflect either generally high copia transcriptional activity or a transition from a “burst” phase to a stabilizing or “cleanup” phase of LTR genome colonization. We therefore compared the number of intact and recently inserted transposable elements with the number of isolated elements (solo:intact) ratio, which reflects the relative rates of proliferation and removal through illegitimate recombination. In general, a higher solo:intact ratio reflects more extensive removal. The solo:intact ratio of copia elements was significantly higher than that of Ty3 elements (copia: mean 5.30, n=1645 families; Ty3, mean 1.86, n=3652 families; t statistic 2.3154 with 417.4 df, p = 0.02107; Test 6 in analysis script), suggesting that even though they are proliferating, they are also being removed more rapidly. For Ty3 elements, age was weakly but significantly positively correlated with distance to genes (i.e., elements closer to genes are both younger and more likely to be removed), while solo:intact ratio was not; neither correlation was significant for copia elements (Table 4; Test 7 in analysis script). This may reflect ongoing proliferation genome-wide: both fragmented and intact copia elements are evenly distributed across the genome, and may be both inserting and being removed too rapidly to leave signatures of differential removal near genes.

**Table 4:** Correlations between family-wide mean age, family-wide mean distance to genes, and family-wide mean solo:intact ratio. All cells indicate Pearson correlations.

| LTR class | Correlation | df | t | p | r |
| --- | --- | --- | --- | --- | --- |
| Ty3 | Age : distance | 2463 | 4.331 | 1.541e-05 | 0.087 |
| Ty3 | Age : distance | 4926 | 1.312 | 0.1896 | 0.019 |
| Copia | Age : solo:intact ratio | 397 | 0.1881 | 0.8509 | 0.009 |
| Copia | Age : solo:intact ratio | 796 | -0.233 | 0.8159 | -0.008 |

Finally, we investigated possible mechanisms for the dynamic balance between proliferation and removal, and whether this balance changes during LTR diversification. If young copias are expanding into typically Ty3-dominated pericentromeric regions, either by proliferating there or by escaping removal, we would expect young copias, particularly in the largest families, to be further from genes. If, in contrast, LTR niches remain consistent, we would not expect a difference in distribution between age classes. We found that the intact elements in the largest families of both copia and Ty3 elements were further away from genes than elements not in the largest families, suggesting that successful families in both superfamilies may persist through remaining far from genes. However, this pattern could also simply result from LTRs near genes being more generally removed, and from more LTRs from the largest families persisting in all locations (Table 5; Test 8 in analysis script).

**Table 5:** intact elements in the largest LTR families are further from genes.

| LTR class | Largest LTR families mean distance (n) | Other LTR families mean distance (n) | df | t | p |
| --- | --- | --- | --- | --- | --- |
| Ty3 | 148.8kb<br>(7616) | 135.2kb<br>(18853) | 13803 | 6.1256 | 9.28e-10 |
| Copia | 138.3kb<br>(22649) | 133.1kb<br>(12459) | 27175 | 2.8754 | 0.004038 |

Within copia, however, age and distance to genes were positively correlated for elements in the largest young families (i.e., older elements were further from genes and younger elements were closer to genes; Pearson correlation with 16500 df, t=4.916, p=8.933e-07, r=0.038), while this correlation was non-significant and negative for young families that were not among the largest (i.e., younger elements were further from genes; Pearson correlation with 1534 df, t=-1.0753, p=0.2824, r=-0.027), and a Z-statistic from Fisher’s r-to-z transformation indicated that the slopes differed significantly (Z=2.46, two-sided p=0.0139; Test 9 in analysis script). We interpret this to mean that large, expanding families are not preferentially inserting further from genes, but are instead inserting in many locations and being removed from genic regions by natural selection. Consistent with this, solo transposable elements (remnants of removal by ectopic recombination) in the largest copia families retained higher similarity to reference sequences when further from genes, suggesting less thorough removal (solo TEs in largest copia families n=11590, mean coverage 0.837; solo TEs in other copia families n=53,060, mean coverage 0.957, t statistic -60.293 with 14275=4 df, p < 2.2e-16; Test 10 in analysis script). This is consistent with selection against insertions near genes, which has been documented in multiple systems, and is likely to be more apparent in copia elements because they are more likely to insert closer to genes (Sultana et al. 2017), and with the role of recombination in TE removal (gene-dense regions of large plant genomes typically have higher rates of recombination; (Kent et al. 2017; Brazier and Glémin 2022).

## Conclusions

The genus *Hesperis* includes the largest genomes in the Brassicaceae, and offers a unique opportunity to understand mechanisms of genome expansion. In our high-quality *H. matronalis* genome, we investigated the timing of bursts of transposable elements, and found evidence that the proliferation of Ty3 elements preceded the proliferation of copia elements, with copia elements being both more intact and younger than the more numerous Ty3 elements. Transposable element profiles were broadly similar across a wide sample of central North American accessions. Both clades diversified unevenly within the *H. matronalis* genome compared to other Brassicaceae, with particularly dramatic expansion in the copia taxa Angela and Ale and the Ty3 taxa Athila, CRM, and Retand. We did not find evidence that insertion sites shifted over the course of LTR bursts; rather, removal appears to be concentrated in gene-dense regions. This research represents a valuable comparative resource for research within and beyond the Brassicaceae, and sheds light on mechanisms of genome evolution.

## Materials and Methods

### Genome sequencing and assembly

Flow cytometry genome size estimates were obtained using *Monstera* or *Clivia* standards from Plant Cytometry Services (Didam, The Netherlands) for 326 samples from 33 populations of *Hesperis matronalis* sampled from Ohio and Michigan (Table S1). An individual from the apparent diploid population RIGA (41.816, -83.824) was selected for genome sequencing.

Sequencing and assembly were performed by Dovetail Genomics (Scotts Valley, CA; see supplementary materials and methods). The genome was sequenced to a depth of between 59x and 65x depending on flow cytometry estimate 16.7 million PacBIO CCS (circular consensus sequencing) reads totaling 235.5gbp. An initial assembly was created using Hifiasm v0.15.4-r347 (Cheng et al. 2021) with default parameters and filtered using blobtools v1.1.1 (Laetsch and Blaxter 2017) and purge_dups v1.2.5 (Guan et al. 2020). Completeness was assessed using the basic universal single copy orthology (BUSCO) gene set (Seppey et al. 2019), version eukaryota_odb10. This initial assembly was then scaffolded using an OmniC ligated library and HiRise scaffolding software.

We performed subsequent analyses using the HiRise scaffolded assembly, processed to remove all contigs below either 1Mb or 50Mb in size (retaining scaffolds 1-46 or 1-10) and simplify scaffold names.

### RNASeq

Floral tissue samples of *Hesperis matronalis* were collected from a single natural population at the Koffler Scientific Reserve at Joker’s Hill (KSR; King City, Ontario, Canada; 44.030, -79.530). Individuals were sampled across the natural floral color variation present at the site, including both purple- and white-flowered morphs. All samples originated from the same population and were photographed in the field, with collections made in late June 2017; corresponding seed collections from putative maternal plants were made in mid-September of the same season.

Total RNA was extracted from leaf and floral tissues and sequenced as paired-end Illumina libraries. Libraries were multiplexed and sequenced to high depth, yielding approximately 57-93 million read pairs per sample with mean Phred quality scores of ∼38. Raw reads were adapter-trimmed and quality-filtered prior to downstream analyses.

### Gene annotation

We generated a genome annotation using BRAKER3 (Gabriel et al. 2024) version 3.03 using as evidence the 20 paired-end RNAseq samples described above and protein models from orthodb version 10 (Kriventseva et al. 2019; Gabriel et al. 2024). We assessed annotation completeness using BUSCO version 5.4.4 (Simão et al. 2015) with the odb10 brassicales database.

### Centromere annotation

We identified putative centromeric satellite repeats using TRASH v. 2.0 (Wlodzimierz et al. 2023), retaining the most common 25 repetitive sequences (Table S6) for downstream analysis. Possible centromere locations were described using a combination of score distributions from TRASH, contig self-alignments generated with MUMmer:nucmer (Marçais et al. 2018), and comparisons using the R package pwalign_1.0.0 (citation) between our satellite sequences and satellites from several species in the *Hesperis* clade (Hloušková et al. 2019).

### TE annotation and comparative genomics

We annotated TEs in the *H. matronalis* genome using EDTA version 2.2 and panEDTA version 0.2 (Ou et al. 2019). In addition, we performed simultaneous de novo TE annotations of 11 genomes across the Brassicaceae (Table 2) and re-annotated all genomes using panEDTA (Ou et al. 2024), adding the TAIR12 *Arabidopsis thaliana* curated TE library (Reiser et al. 2024). We limited our downstream analyses of TEs to Long Terminal Repeats (LTRs) in families that were represented by at least one full-length copy in at least one of the genomes explored, retaining 13390 distinct LTRs in our library. We classified our LTRs using TEsorter (Zhang et al. 2022), using a protein domain similarity in the first pass and a 70-30-80 rule to classify initially unclassified LTRs in the second pass. We then aligned identified protein sequences using mafft (Katoh et al. 2009) and constructed phylogenies in IQ-TREE (Minh et al. 2020).

We manually curated our TESorter LTRs, removing duplicate IDs with different domains, duplicate IDs with less complete classification information, and one ID with with contradictory classification information (panTE_00007343, which is classified variously as copia, Ty3, unknown LTR, pararetrovirus, target site deletion). We then curated the panEDTA output used for broad-scale patterns as follows: when different copies of an LTR had conflicting annotation but the LTR was present in TESorter, we applied the classification from TESorter, and when LTRs had two conflicting annotations but one was “LTR/Unknown,” the other was applied. Based on these criteria, we retained a final list of 10250 LTR families.

We used custom R scripts to quantify LTR burden in *H. matronalis* and other species. For intact long terminal repeat retrotransposons, we estimated insertion age using the formula "T = K/2*r with T = K/2 × r,” where T = time of divergence, K = divergence and r = substitution rate (Jedlicka et al. 2020). For substitution rate, we used the *Arabidopsis* estimate of 1.5 × 10^–8^ (Koch et al. 2000).

We compared synteny across genomes using GENESPACE (Lovell et al. 2022).

### TE quantification from field samples

In addition to our *H. matronalis* genome assembly, we used the TERAD pipeline (Chak and Rubenstein 2019) to quantify approximate TE composition from RADseq sequencing from 480 individuals from 47 wild populations with EcoR1 and MSP1 (Johnson et al. 2025). TERAD clusters RADseq reads and compares clusters to repeat libraries; we pooled all samples from each population into a single analysis for every site.

## Data Availability

All analysis code will be available at a GitHub repository for the manuscript. The genome assembly, annotations, and RNASeq will be deposited with EMBL/ENA. RADSeq sequencing reads from (Johnson et al. 2025) will be deposited at the Sequence Read Archive (SRA).

