## Supplemental Methods and Figures for "The Dynamics of Long Terminal Repeat Proliferation in the *Hesperis matronalis* Genome"

### Supplementary Methods and Results

**Genome sequencing, assembly, and scaffolding of *Hesperis matronalis***
High-molecular-weight genomic DNA from *Hesperis matronalis* was sequenced and assembled using a hybrid long-read and proximity-ligation approach. DNA was submitted to Dovetail Genomics LLC (Scotts Valley, CA, USA) for PacBio HiFi sequencing, de novo assembly, Omni-C proximity ligation library construction, deep sequencing, and HiRise scaffolding.

PacBio HiFi sequencing was performed on four SMRT Cells, generating high-fidelity long reads that were assembled de novo using hifiasm, producing a haplotype-resolved primary assembly. The resulting contigs were subsequently scaffolded using Omni-C chromatin conformation data.

235.5 gigabase-pairs of PacBio CCS reads were used as an input to Hifiasm1 v0.15.4-r347 with default parameters. Blast results of the Hifiasm output assembly (hifiasm.p_ctg.fa) against the nt database were used as input for blobtools2 v1.1.1 and scaffolds identified as possible contamination were removed from the assembly (filtered.asm.cns.fa). Finally, purge_dups3 v1.2.5 was used to remove haplotigs and contig overlaps (purged.fa). See below for all software versions.

For Omni-C library preparation, chromatin was fixed in nuclei using formaldehyde, extracted, and digested with DNase I. DNA ends were repaired, and biotinylated bridge adapters were ligated prior to proximity ligation. Crosslinks were reversed, DNA was purified, and biotinylated fragments were enriched using streptavidin beads following library construction with NEBNext Ultra enzymes and Illumina-compatible adapters. The final library was sequenced on an Illumina HiSeq X platform to approximately 30× coverage. Reads with mapping quality ≥50 were retained for scaffolding.

Hi-C–based scaffolding was performed using the HiRise pipeline (Dovetail Genomics), which aligns Omni-C reads (MQ>50 used) to the draft assembly using BWA and models genomic distance distributions between read pairs to identify misjoins and evaluate candidate scaffold joins. Iterative correction was used to break putative misassemblies and introduce high-confidence joins.

<u>Assembly statistics and improvement after scaffolding</u>
HiRise scaffolding substantially improved assembly contiguity while maintaining overall assembly size. The input hifiasm assembly had a total length of 3,871,023,182 bp, which was essentially unchanged after scaffolding (3,871,040,282 bp). HiRise introduced 158 scaffold joins and 3 breaks using 54,540,871 read pairs.

Contiguity metrics improved markedly following scaffolding. Scaffold N50 increased from 33.57 Mb in the input assembly to 405.65 Mb in the HiRise assembly, while L50 decreased from 36 to 5. N90 increased from 3.63 Mb to 9.80 Mb, with L90 decreasing from 146 to 15. The largest scaffold increased from 116.71 Mb in the input assembly to 446.71 Mb in the final assembly. The total number of scaffolds decreased slightly from 6,411 to 6,243, all exceeding 1 kb in length. The final assembly contained 158 scaffold gaps, corresponding to 0.44 Ns per 100 kb.

Assembly completeness

Assembly completeness was assessed using BUSCO v4.0.5 with the eukaryota_odb10 dataset (255 conserved orthologs). The input assembly contained 254 complete BUSCOs (99.61%), including 133 single-copy and 121 duplicated genes, with 1 fragmented and 0 missing BUSCOs. The final HiRise assembly contained 253 complete BUSCOs (99.22%), including 136 single-copy and 117 duplicated BUSCOs, with 1 fragmented and 1 missing BUSCO.

**BUSCO completeness assessment**

| Assembly | Complete BUSCOs | Single copy | Duplicated | Fragmented | Missing | BUSCO groups |
|---|---|---|---|---|---|---|
| Input assembly | 254/255 (99.61%) | 133 | 121 | 1 | 0 | 255 |
| HiRise assembly | 253/255 (99.22%) | 136 | 117 | 1 | 1 | 255 |

#### Dovetail software versions

| *Package* | *Version* |
|---|---|
| awscli | 1.20.0 |
| bioawk | 1.0 |
| blas | 1.0 |
| blast | 2.9.0 |
| blobtools | 1.1.1 |
| minimap2 | 2.21 |
| numpy | 1.19.1 |
| pandas | 1.1.3 |
| pip | 20.2.4 |
| pyqt | 5.9.2 |
| pysam | 0.15.4 |
| python | 3.7.6 |
| qt | 5.9.7 |
| samtools | 1.9 |

wtdbg 2.5

### Supplementary Figures

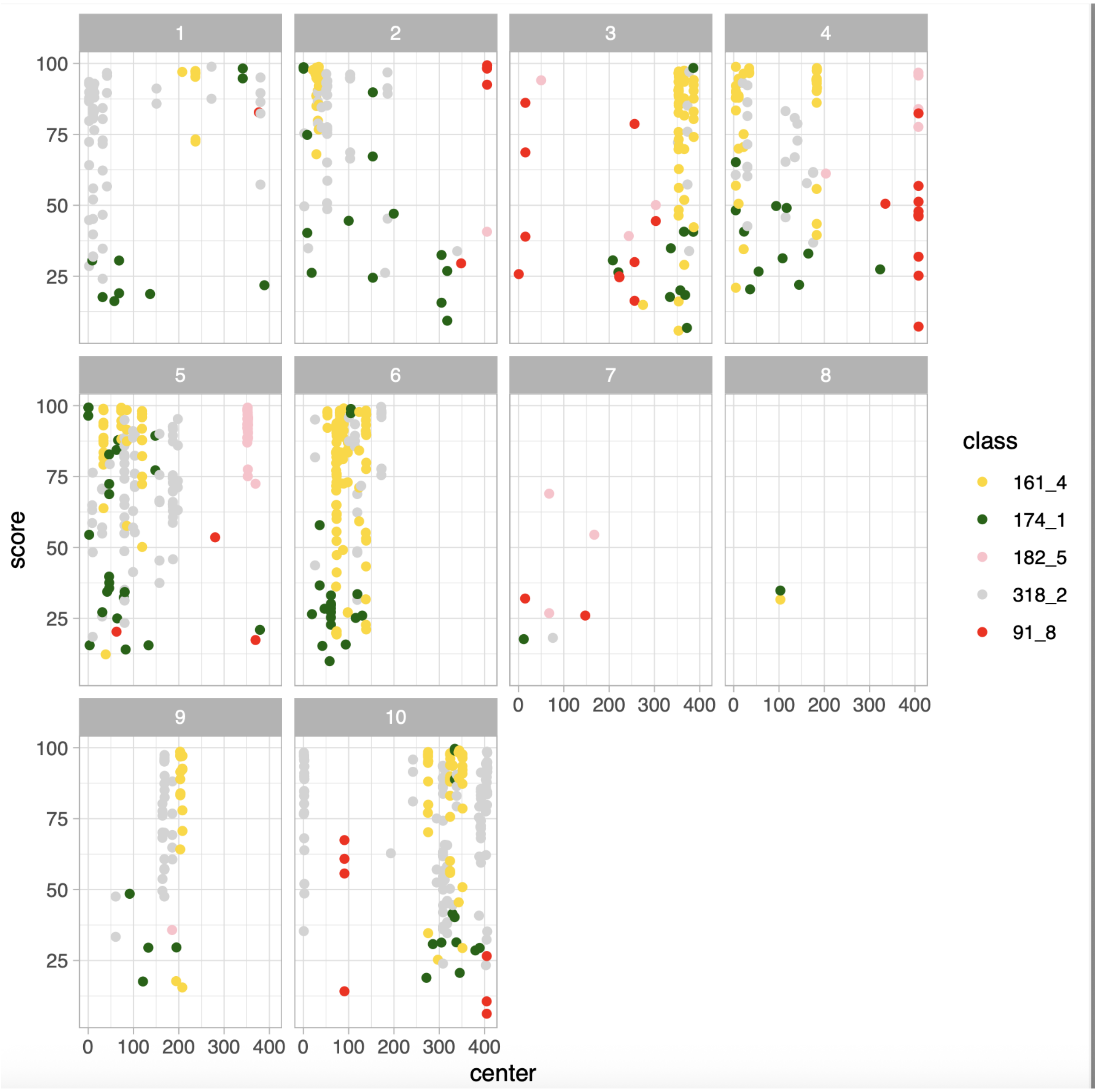


Figure S1. Positions of satellites on major scaffolds.
